# Selective elimination of CD169^+^ macrophages in lymph nodes invaded by breast cancers

**DOI:** 10.1101/2023.08.02.551659

**Authors:** Yurina Maeshima, Tatsuki R. Kataoka, Alexis Vandenbon, Masahiro Hirata, Yasuhide Takeuchi, Yutaka Suzuki, Yukiko Fukui, Yumiko Ibi, Hironori Haga, Satoshi Morita, Masakazu Toi, Shinpei Kawaoka, Kosuke Kawaguchi

**Affiliations:** Department of Breast Surgery, Kyoto University Hospital, Graduate School of Medicine, Shogoin Sakyo-ku, Kyoto 606-8507, Japan; Inter-Organ Communication Research Team, Institute for Frontier Life and Medical Sciences, Kyoto University, Sakyo-ku, Kyoto 606-8507, Japan; Department of Pathology, Iwate Medical University, Yahaba-cho, Shiwa-gun, Iwate Prefecture 028-3694, Japan; Laboratory of Tissue Homeostasis, Institute for Life and Medical Sciences, Kyoto University, Sakyo-ku, Kyoto 606-8507, Japan; Institute for Liberal Arts and Sciences, Kyoto University, Kyoto 606-8507, Japan; Department of Diagnostic Pathology, Kyoto University, Shogoin Sakyo-ku, Kyoto 606-8507, Japan; Graduate School of Frontier Science, The University of Tokyo, Chiba 277-8562, Japan; Department of Biomedical Statistics and Bioinformatics, Kyoto University Graduate School of Medicine, Shogoin Sakyo-ku, Kyoto 606-8507, Japan; Director, Tokyo Metropolitan Cancer and Infectious Disease Center, Komagome Hospital, Honkomagome, Bunkyo-ku, Tokyo, 113-8677, Japan; Department of Integrative Bioanalytics, Institute of Development, Aging and Cancer, Tohoku University, Sendai 980-8575, Japan

## Abstract

Breast cancer cells suppress the host immune system to efficiently invade the lymph nodes; however, the underlying mechanism remains incompletely understood. Here, we report that metastasized breast cancer cells selectively eliminate CD169^+^ lymph node sinus macrophages, an initiator of anti-cancer immunity, from the lymph nodes. The comparison between paired lymph nodes with and without metastasis from the same patients demonstrated that the number of CD169^+^ macrophages was reduced in metastatic lymph nodes, whereas the numbers of other major immune cell types were unaltered. We also detected the infiltration of CD169^+^ macrophages into metastasized cancer tissues depending on sections, suggesting that CD169^+^ macrophages were gradually eliminated after anti-cancer reactions. Furthermore, CD169^+^ macrophage elimination was prevalent in major breast cancer subtypes and correlated with breast cancer staging. Collectively, we propose that metastasized breast cancer cells dispel CD169^+^ macrophages from lymph nodes in a phased manner, disabling a critical step of anti-cancer immunity.

## Introduction

Cancers affect host cells in various ways to ensure their survival and evade the host immune system^1–4^. Cancer cells directly contact host immune cells via cell surface molecules, inhibiting immune cell activity^5, 6^. Cancer cells also secrete soluble factors, including cytokines, that influence nearby cells^7, 8^. Cancer-derived factors can have distant effects when they enter the blood and lymph streams^9–11^. Such direct, proximal, and distal interactions between cancer and host cells eventually disrupt host homeostasis^4, 9, 12–15^. These cancer–host interactions are even more complex in the presence of more than two cancer tissues (e.g., primary and metastasized cancer tissues) within the same body^16^.

The lymph nodes represent the main site of action of such complex cancer–host interactions^17^. The interaction between cancer cells and lymph nodes is of particular interest because metastasis to the lymph nodes requires the efficient suppression of host immune cells^17^. Previous studies demonstrate immune cell suppression in the lymph nodes of cancer patients, as exemplified by the increase in the regulatory T cell (T_reg_) population^18, 19^. Given the immunosuppressive function of T_reg_ cells, this is a reasonable mode of immune suppression by cancer cells^18, 19^. However, it remains incompletely understood whether other immune cell types are affected in the lymph nodes of cancer-bearing organisms. Moreover, distinguishing the direct, proximal, and long-range effects of primary and secondary cancer tissues on host immune cells in the lymph nodes is challenging. Addressing these issues requires a comprehensive dissection of cancer–host interactions within the lymph nodes, which is critical for precisely understanding the mechanisms by which cancer cells colonize this immune cell-rich organ in the body.

CD169^+^ macrophages are a unique type of resident macrophages in the lymphoid organs that contribute to anti-cancer immunity^20–23^. CD169^+^ macrophages phagocytose dead cancer cells and present cancer-derived antigens to CD8^+^ T cells^20^. Animal studies have revealed the significant contribution of CD169^+^ macrophages to the suppression of cancer growth and metastasis *in vivo*^21^. Clinical studies have also supported the importance of CD169^+^ macrophages; the number of CD169^+^ macrophages in the lymph nodes is positively correlated with the prognosis of patients with various cancer types^24–29^. However, the functional significance of CD169^+^ macrophages in patients with cancer remains unclear. Are CD169^+^ macrophages major targets of cancer cells in patients? If yes, when do cancer cells start to affect this cell type? These questions have been unanswered particularly in patients.

In the current study, we aimed to comprehensively characterize the effects of breast cancers on immune cells in the lymph nodes. We analyzed non-metastatic and metastatic lymph node samples from patients with breast cancer with lymph node metastasis using multi-scale transcriptome analyses. The obtained datasets were used to capture the direct and proximal effects of metastasized cancer cells on host immune cells in a relatively unbiased manner, providing new insight into CD169^+^ macrophages as targets of metastasized breast cancer cells in the lymph nodes. This study thus deepens our understanding of immunosuppression by metastasized breast cancer cells in the lymph nodes, establishing CD169^+^ macrophages as crucial therapeutic targets.

## Results

### Searching for lymph node genes affected by breast cancer metastasis

To determine the direct and proximal effects of metastatic breast cancer cells on host cells in the lymph nodes, we collected 20 laser micro-dissected sections from 17 lymph nodes of six patients with breast cancer who had lymph node metastasis without clinical evidence of distant organ metastasis (i.e., stages II–III; Fig. 1a and Table 1). Four patients had breast cancers expressing hormone receptor proteins (luminal-type breast cancer), and breast cancers of two patients did not express hormone receptor proteins and did not exhibit *HER2* amplification (triple-negative breast cancer subtype). Each patient had at least one non-metastatic and one metastatic lymph node, allowing direct comparison between paired non-metastatic and metastatic lymph nodes in the same patient (Table 1). Given the long-range effects of primary breast cancers via the lymph^30–34^, this comparison was considered to be useful for identifying the direct and proximal effects of cancer cells on host immune cells in the lymph nodes. In four cases, the patient had more than two non-metastatic and/or metastatic lymph nodes (Table 1). In these cases, we averaged the obtained gene expression data to normalize variations among lymph nodes from the same patient. As a result, we generated 12 gene expression matrices (six non-metastatic and six metastatic lymph nodes) (Extended Data Table 1).

**Fig. 1.**
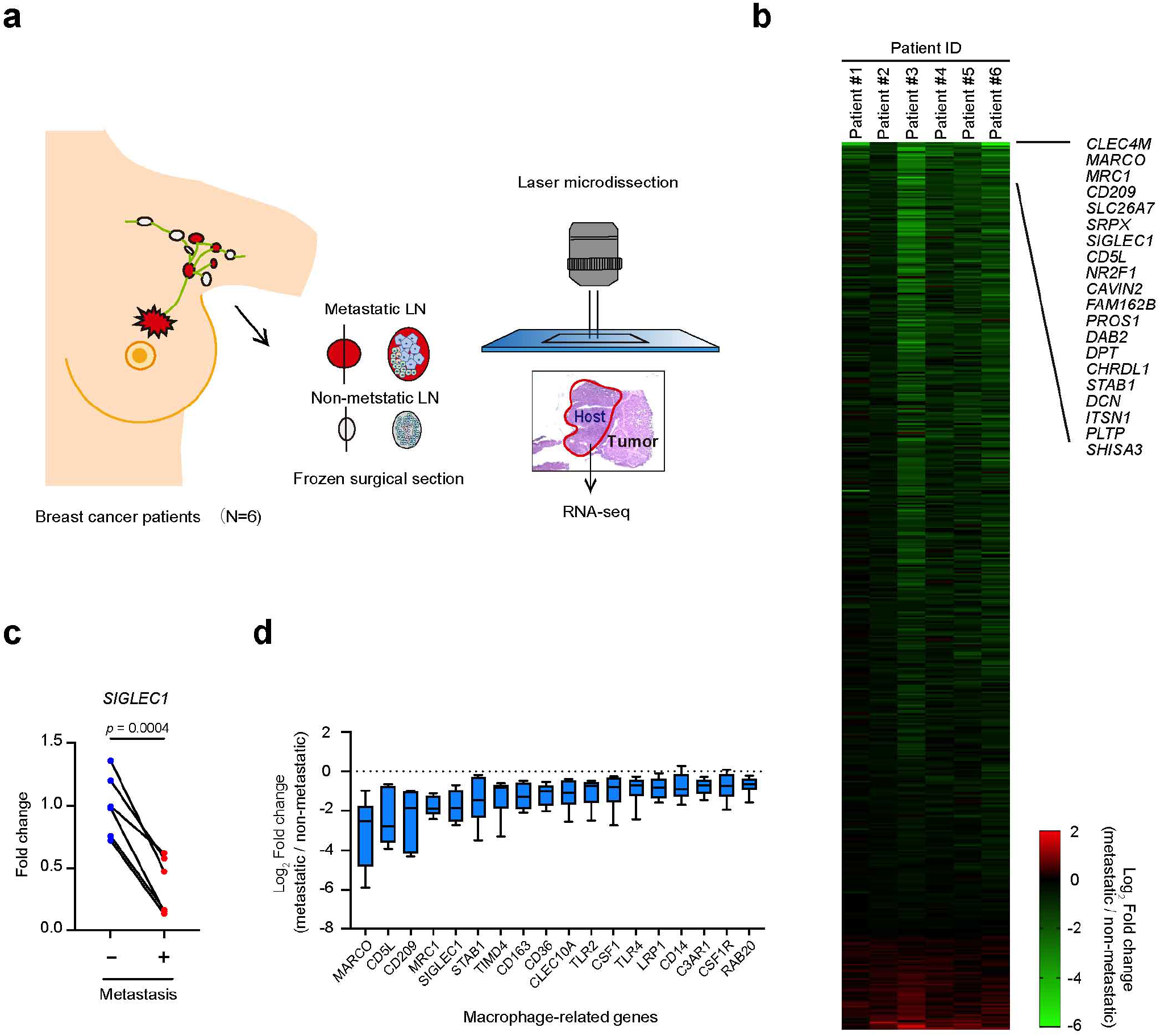
Reduced expression of macrophage-related genes in lymph nodes with breast cancer metastasis. **a.** Experimental workflow. **b.** Heatmap of genes differentially expressed in the metastatic lymph nodes. Gene expression in metastatic lymph nodes was normalized to that in non-metastatic lymph nodes for each patient (log_2_ fold change). The top 20 most downregulated genes are highlighted; *n* = 6. **c**. Expression of *SIGLEC1* (RNA-seq). The average fold-change data normalized to non-metastatic lymph nodes are presented. The *p-*value was calculated using the paired two-tailed Student *t*-test; *n* = 6. **d.** Expression of macrophage-related genes in the lymph nodes (RNA-seq). Gene expression in the metastatic lymph nodes was normalized to that in non-metastatic lymph nodes for each patient (log_2_ fold change); *n* = 6.

**Table 1.** Patient characteristics and information of the collected lymph node samples.

|  | Patient 1 | Patient 2 | Patient 3 | Patient 4 | Patient 5 | Patient 6 |
| --- | --- | --- | --- | --- | --- | --- |
| Age | 54 | 41 | 45 | 76 | 69 | 68 |
| Pathological T category | 4b | 2 | 2 | 2 | 2 | 2 |
| Pathological N category | 2a | 3a | 2a | 1a | 1a | 3a |
| N status | 4/14 | 12/20 | 4/12 | 2/13 | 2/15 | 11/16 |
| Tumor subtype | Luminal | Luminal | Luminal | TNBC | Luminal | TNBC |
| Ki67 (%) | 33.2 | 22 | 98.6 | 65 | 62 | 80.2 |
| No. of collected lymph nodes |  |  |  |  |  |  |
| metastatic | 1 | 2 | 1 | 1 | 1 | 2 |
| non-metastatic | 1 | 1 | 1 | 2 | 2 | 2 |
| No. of laser micro-dissected sections |  |  |  |  |  |  |
| metastatic | 1 | 2 | 1 | 4 | 1 | 2 |
| non-metastatic | 1 | 1 | 1 | 2 | 2 | 2 |

Abbreviation: TNBC, triple-negative breast cancer

A total of 489 differentially expressed genes (*P* < 0.05) were identified between the paired non-metastatic and metastatic lymph nodes. Notably, most of the affected genes were downregulated in the metastatic samples (Fig. 1b). These differentially expressed genes included gene expression changes that were reported previously^32^. For example, the expression levels of *Chemokine (C-C-motif) Ligand 21* (*CCL21*) and *Interleukin-7* (*IL-7*) were reduced in the metastatic lymph nodes (Extended Data Fig. 1a, b). These two genes are representative of cancer-dependent lymph node reprogramming^32^, which is considered to occur via long-range mechanisms. Thus, our data suggested that cancer cells further remodel the structure of the lymph nodes upon arrival.

A closer examination of the heatmap of differential gene expression revealed that genes expressed in macrophages were preferentially affected by the presence of lymph node metastasis. Notably, the expression level of sialic acid binding IG-like lectin1 (*SIGLEC1*; also known as *CD169*), the canonical marker of CD169^+^ macrophages^23, 35^, was severely reduced in lymph nodes harboring metastasis (Fig. 1c, d). In addition, reductions in macrophage receptor with collagenous structures (*MARCO*) and apoptosis inhibitor of macrophage (*AIM;* also known as *CD5L*) in metastatic lymph nodes were also notable (Fig. 1d and Extended Data Fig. 1c). Collectively, our transcriptome analyses indicated that the direct and proximal effects of metastatic breast cancer suppress a set of genes expressed in macrophages.

### Spatial transcriptomics reveals the reduction of lymph node macrophages

We next investigated whether the identified changes in host genes reflected the quantity and quality changes in immune cell populations using spatial transcriptome analysis. We selected two sets of non-metastatic and metastatic lymph node pairs from two patients (patients #4 and #6) to obtain four slices. After a quality check and batch-effect correction, 12,625 spots were obtained for further analysis. The datasets were subjected to dimensionality reduction using principal component analysis and Uniform Manifold Approximation and Projection (UMAP), revealing 10 clusters in the lymph nodes (Fig. 2a). These clusters were annotated according to known marker genes (Extended Data Table 2). We also calculated “module scores” (averaged expression levels of genes involved in the same biological processes; see Methods) using Seurat for sets of genes associated with 3152 Gene Ontology (GO) terms for annotation. These analyses identified B cells, T cells, macrophages, plasmacytoid dendritic cells (pDCs), endothelial cells, and cancer cells in our lymph node datasets (Fig. 2a, b and Extended Data Fig. 2a–f).

**Fig. 2.**
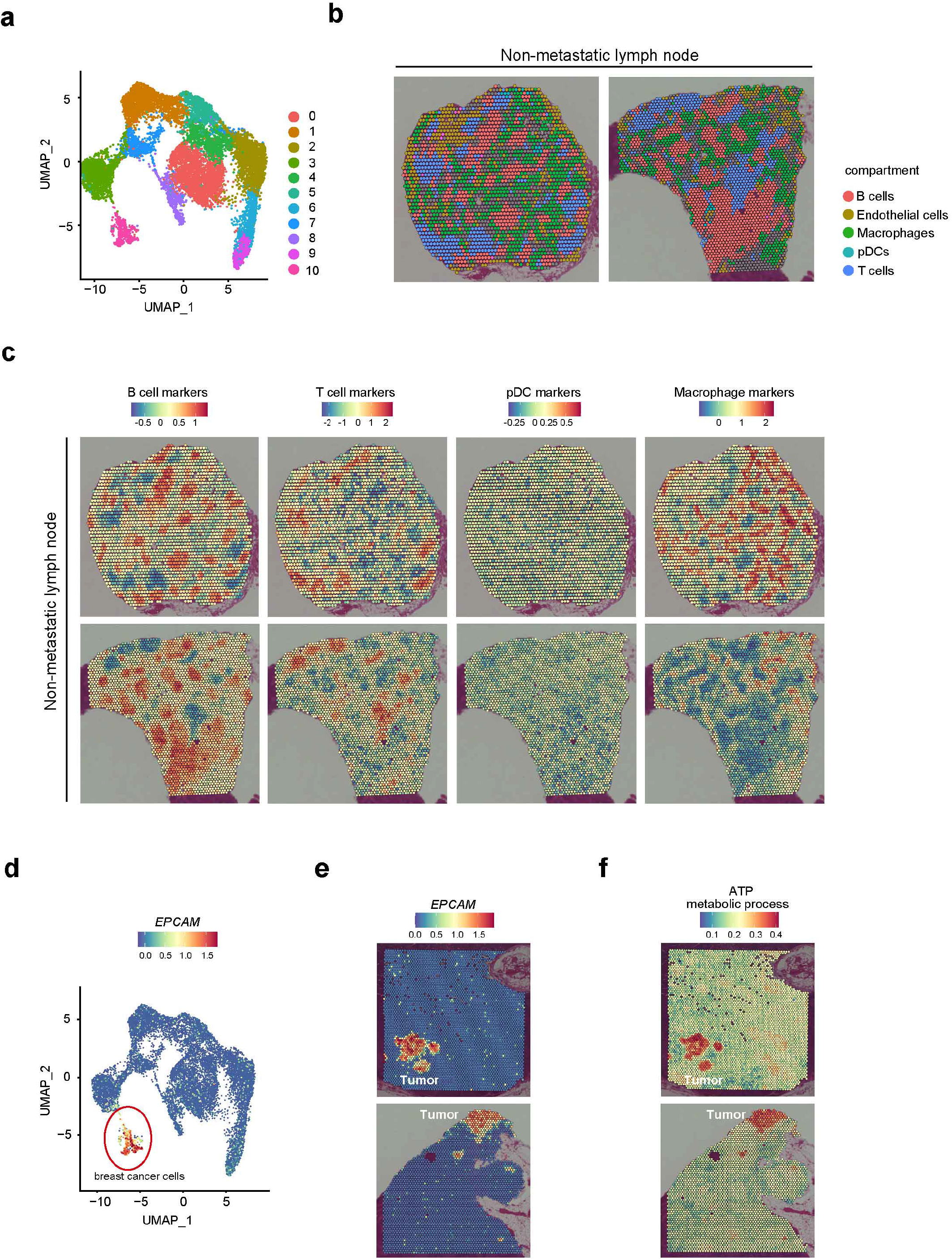
Spatial transcriptomics landscape of lymph nodes of breast cancer patients. **a.** UMAP plot of VISIUM spots of non-metastatic and metastatic lymph nodes. Two pairs of non-metastatic and metastatic lymph nodes from patients #4 and #6 are presented. **b.** Compartmentalization of lymph node cells. Each cell type was identified using a specific set of marker genes listed in Extended Data Table 2 (also see Extended Data Fig. 2). **c.** Gene Ontology (GO)-based annotation of lymph node cell types. VISIUM spots enriched for B cells, T cells, plasmacytoid dendritic cells (pDCs), and macrophage markers are presented. **d.** Expression of *EPCAM* in UMAP plots. *EPCAM*-high spots are highlighted as metastatic breast cancer cells. **e.** Expression of *EPCAM* in the VISIUM slices from metastatic lymph nodes. *EPCAM*-high spots correspond to histologically annotated tumor areas. **f.** Module scores for “ATP metabolic process” in the VISIUM slices from metastatic lymph nodes.

Consistent with the previously established structure of the lymph nodes^36^, these different immune cell types were observed to be compartmentalized rather than uniformly distributed in the lymph nodes (Fig. 2b, c). Using two slices from non-metastatic lymph nodes as examples, B cell- and T cell-enriched spots demonstrated distinct patterns of localization in an almost mutually exclusive manner (Fig. 2c). Although our spatial transcriptomic data were not obtained at the single-cell level, based on these results, each spot was expected to be enriched in specific cell types. Macrophage and pDC spots also exhibited a unique distribution and rarely overlapped with B cell- and T cell-enriched spots (Fig. 2c).

Our analyses successfully captured metastasized breast cancer cells (Fig. 2d; cluster 10). The cluster corresponding to the metastasized breast cancer cells highly expressed *EPCAM*, an epithelial marker gene (Fig. 2e). These cancer cells were enriched for a GO term “ATP metabolic process,” suggestive of active energy production and usage (Fig. 2f). Spots surrounding cancer cells, which expressed *NNMT* at the high level, were expected to be metastasis-associated stroma as mentioned previously in ovarian cancers (Extended Data Fig. 2g)^37^. Furthermore, the metastasized lymph nodes had more than two foci of breast cancer cells within the sections (Fig. 2e). Thus, these lymph nodes appeared in the process of being occupied by metastasized breast cancer cells. Although similar “cancer spots” were also detected in the non-metastatic lymph nodes, unlike those observed in the metastatic lymph nodes, these spots were sparsely distributed (Extended Data Fig.2h), and they were not histologically considered to be metastasized cancer cells.

Comparing the non-metastatic and metastatic lymph nodes demonstrated that macrophages (cluster 1) represented the most strongly reduced population in breast cancer metastasis (Fig. 3a). The number of spots corresponding to cluster 1 was reduced by more than two-fold in metastatic lymph nodes compared with that in non-metastatic lymph nodes. This observation aligned with our bulk transcriptome data, demonstrating a decrease in a series of macrophage markers (Fig. 1b–d). Notably, *SIGLEC1* was abundantly expressed in this macrophage cluster (Fig. 3b). Furthermore, spots with high expression of *SIGLEC1* were localized in the lymph node sinuses (Fig. 3c), suggesting that these spots contained CD169^+^ lymph node sinus macrophages. Visualization of whole macrophage-enriched spots and *SIGLEC1* (*CD169*)-positive spots in the four slices demonstrated a severe reduction in *SIGLEC1*-positive spots in metastatic lymph nodes (Fig. 3d,e). We detected almost no *SIGLEC1*-positive spots near cancer tissues. In contrast, other cell types (e.g., B and T cells) were not strongly affected by breast cancer metastasis in these sections (Extended Data Fig. 3). These results suggested that macrophages, including CD169^+^ macrophages, in the lymph nodes are selectively affected by metastatic breast cancer cells.

**Fig. 3.**
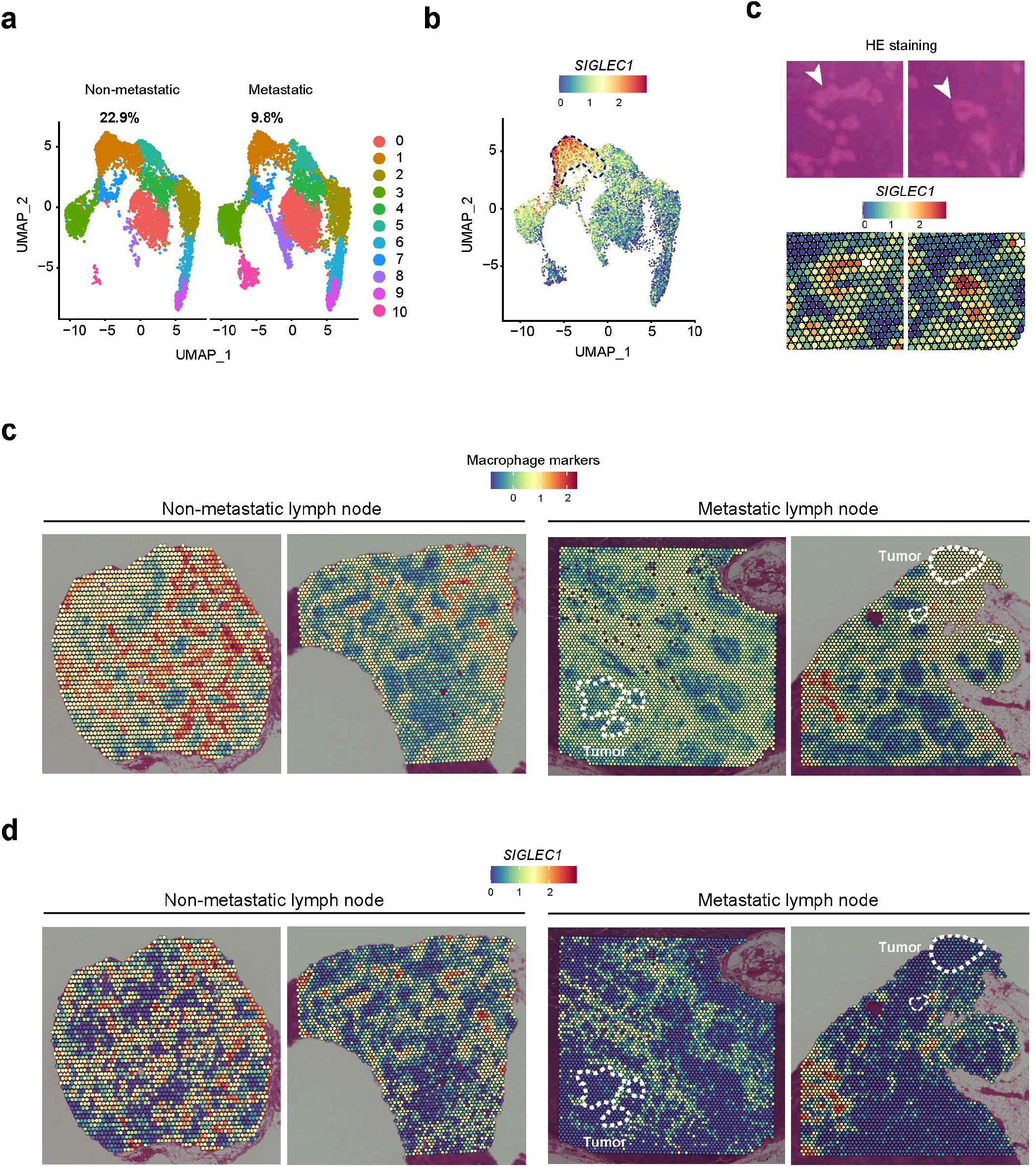
Spatial transcriptomics reveals macrophage reduction in metastatic lymph nodes. **a.** UMAP plots of non-metastatic and metastatic lymph nodes. The fraction of macrophage-enriched spots is presented as the percentage. **b.** Expression of *SIGLEC1* in the UMAP plot. Macrophage-rich spots (Cluster 1) are highlighted. **c**. Localization of *SIGLEC1*-high spots in the lymph nodes. Upper panel: hematoxylin-eosin (HE) staining; lower panel: expression of *SIGLEC1* in VISIUM slices. Arrowheads indicate lymph node sinuses. **d.** Macrophage-rich spots in non-metastatic and metastatic lymph nodes. Metastatic breast cancer cells are outlined. **e.** Expression of *SIGLEC1* in non-metastatic and metastatic lymph nodes. Metastatic breast cancer cells are outlined.

### CD169^+^ macrophages are select targets of metastasized breast cancers

We further tested if CD169^+^ macrophages are selectively reduced in metastatic lymph nodes using the Hyperion imaging system^38^, with which we can visualize cells of interest at the single-cell resolution. We used anti-CD20 antibody for B cells, anti-CD4 antibody for CD4^+^ T cells, anti-CD8 antibody for CD8^+^ T cells, and anti-FOXP3 antibody for T_reg_ cells, anti-CD68 antibody for total macrophages, anti-CD169 (SIGLEC1) antibody for CD169^+^ macrophages, anti-CD11c antibody for CD11c^+^cells (e.g., dendritic cells), anti-α-smooth muscle actin (SMA) for myoepithelial cells, and anti-pan-cytokeratin for cancer cells (Fig. 4a and Table S3: the list of antibodies used) against the same set of samples we used for spatial transcriptomics, enabling direct comparison. We selected 3 regions of interest (ROI) and counted the number of these cell types in each slice (Extended Data Fig. 4).

**Fig. 4.**
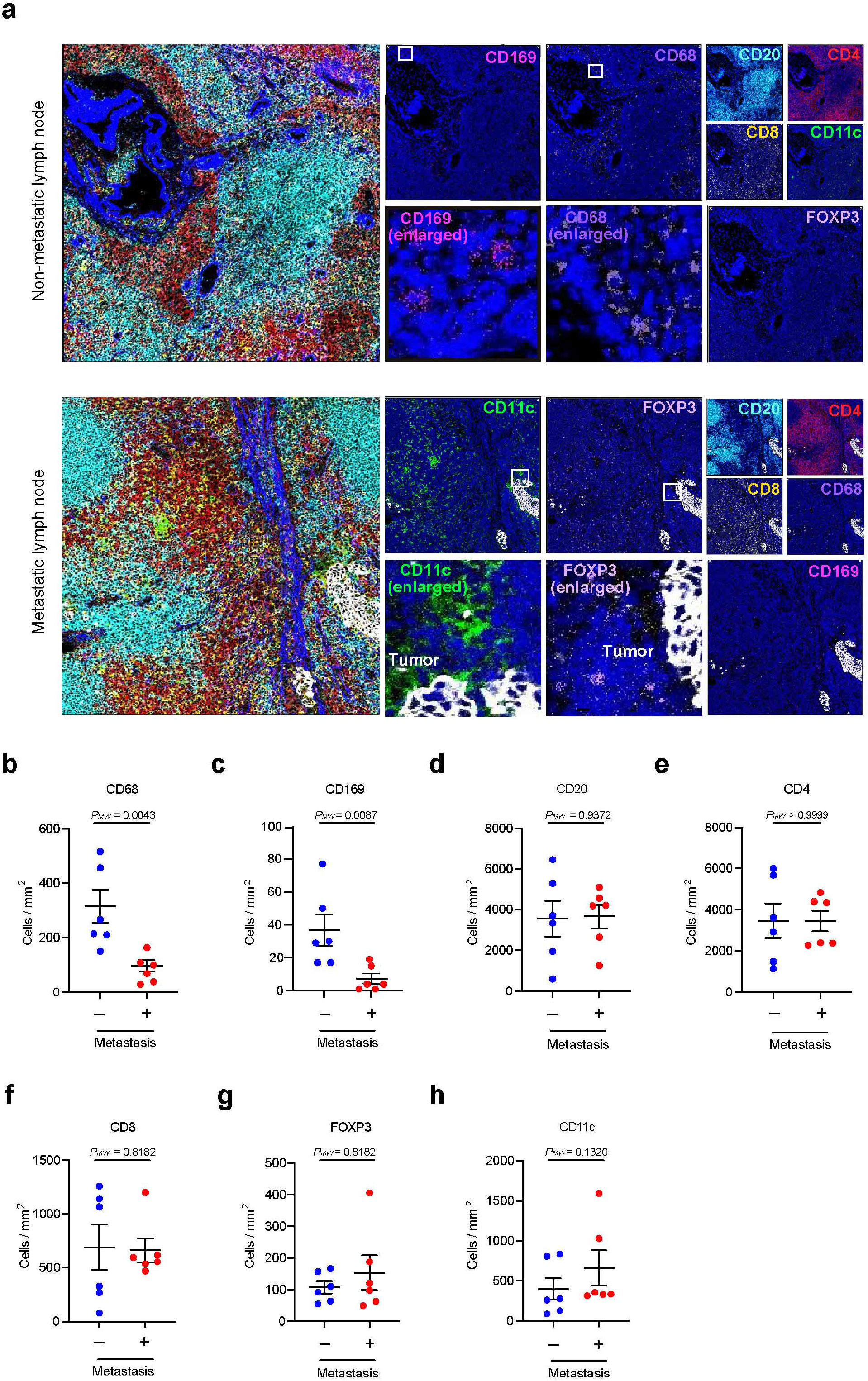
CD169^+^ macrophages are select targets of metastasized breast cancers. **a.** Representative imaging mass cytometry images of non-metastatic and metastatic lymph nodes from patient #6. Displayed channels are as follows: CD20 (B cells; cyan), CD4 (CD4^+^ T cells; red), CD8 (CD8^+^ T cells; yellow), FOXP3 (T_reg_ cells; pale purple), CD68 (macrophages; purple), CD169 (CD169^+^ macrophages; pink), CD11c (CD11c^+^ cells; green), pan-cytokeratin (cancer cells; white), DNA intercalator (nucleus; blue), and anti-α-smooth muscle actin (SMA) (myoepithelial cells; blue). DNA intercalators are displayed in each panel. The SMA staining is depicted in the merged images. Cell phenotypes were determined using HALO software. **b.** The density of CD68^+^ macrophages in non-metastatic and metastatic lymph nodes. **c.** The density of CD169^+^ macrophages in non-metastatic and metastatic lymph nodes. **d**. The density of CD20^+^ B cells in non-metastatic and metastatic lymph nodes. **e.** The density of CD4^+^ T cells in non-metastatic and metastatic lymph nodes. **f.** The density of CD8^+^ T cells in non-metastatic and metastatic lymph nodes. **g.** The density of FOXP3^+^ T_reg_ cells in non-metastatic and metastatic lymph nodes. **h.** The density of CD11c^+^ cells in non-metastatic and metastatic lymph nodes. **(b–h)**. Data were calculated from three regions of interest (ROIs) per section using HALO software (six ROIs per group). Data are presented as the mean ± SEM. The *p* values were calculated with the two-tailed Mann–Whitney *U* test.

As displayed in Fig. 4a, distinct immune cell types were successfully visualized. The compartmentalized localization of these cell types was consistent with the spatial transcriptomic analysis (Fig. 2-3). Importantly, our data demonstrated that the numbers of total macrophages (Fig. 4b) and CD169^+^ macrophages (Fig. 4c) were smaller in the metastatic lymph nodes than in the non-metastatic lymph nodes, suggesting that cancer cells uprooted CD169^+^ macrophages from the lymph nodes.

In contrast to macrophages, the numbers of B, T, T_reg_, and CD11c^+^ cells were comparable in non-metastatic and metastatic lymph nodes (Fig. 4d–h). Furthermore, T_reg_ cells appeared enriched around metastasized cancer tissues, supporting previous findings of the T_reg_-dependent suppression of anti-cancer immunity (Fig. 4a)^18, 19^. Given the unaltered number of T_reg_ cells in metastasized lymph nodes, these results indicate the migration of T_reg_ cells toward cancer tissues (Fig. 4g). We also observed that CD11c^+^ cells were enriched around the cancer tissues (Fig. 4a).

Collectively, we concluded that metastasized breast cancer cells selectively eliminated CD169^+^ macrophages from metastasized lymph nodes, probably suppressing the earlier step of anti-cancer immunity of the host. The examined patients had breast cancers with lymph node metastasis but without clinical evidence of distant organ metastasis (i.e., stage II to III). They had normal levels (i.e., the range of healthy subjects) of albumin and C-reactive protein (CRP), which are used to know the patient’s systemic status (Extended Data Table 4)^39, 40^. Together with relatively earlier breast cancer stages of the examined patients, their systemic statuses were not considered terminal. Based on these results, we speculate that cancer-dependent suppression of CD169^+^ macrophages precedes other cancer-dependent abnormalities of the host, such as systemic inflammation.

### The elimination of CD169^+^ macrophages is a generalizable event in breast cancer

As described earlier, CD169^+^ macrophages play a critical role in anti-cancer immunity^20–23^. In this sense, it is unlikely that CD169^+^ macrophages are suppressed from the beginning without participating in anti-cancer immunity. It is more likely that CD169^+^ macrophages are overwhelmed by cancer cells after some fights, consequently being eliminated from the lymph nodes. In other words, there must be a stepwise mechanism of cancer-dependent suppression of CD169^+^ macrophages. Related to these assumptions, it is of note that our analyses investigated only snapshots of cancer-dependent elimination of CD169^+^ macrophages. In addition, our spatial analysis was limited to a small number of patients. These situations were not ideal for examining such a phased mechanism of elimination of CD169^+^ macrophages. Moreover, the limited number of samples prevented us from generalizing our findings from spatial analyses. To address these issues, we analyzed an additional 315 non-metastatic lymph nodes and 159 metastatic lymph nodes from 58 patients with breast cancer (Extended Data Table 5).

These experiments revealed several crucial findings. First, we confirmed that the number of CD169^+^ macrophages was reduced in the metastatic lymph nodes compared with that in the non-metastatic lymph nodes (Fig. 5a). Thirty-seven of the 159 metastatic lymph nodes revealed no detectable CD169^+^ macrophages, suggesting complete elimination of CD169^+^ macrophages in these samples. Most CD169^+^ macrophages existed in the lymph node sinuses, validating the important nature of this particular macrophage (Fig. 5b). We also noted the consistent reduction of CD169^+^ macrophages in three major breast cancer subtypes (luminal, HER2-positive, and triple-negative breast cancers) (Fig. 5c). These results indicated that the elimination of CD169^+^ macrophages is a general phenomenon in patients with breast cancer.

**Fig. 5.**
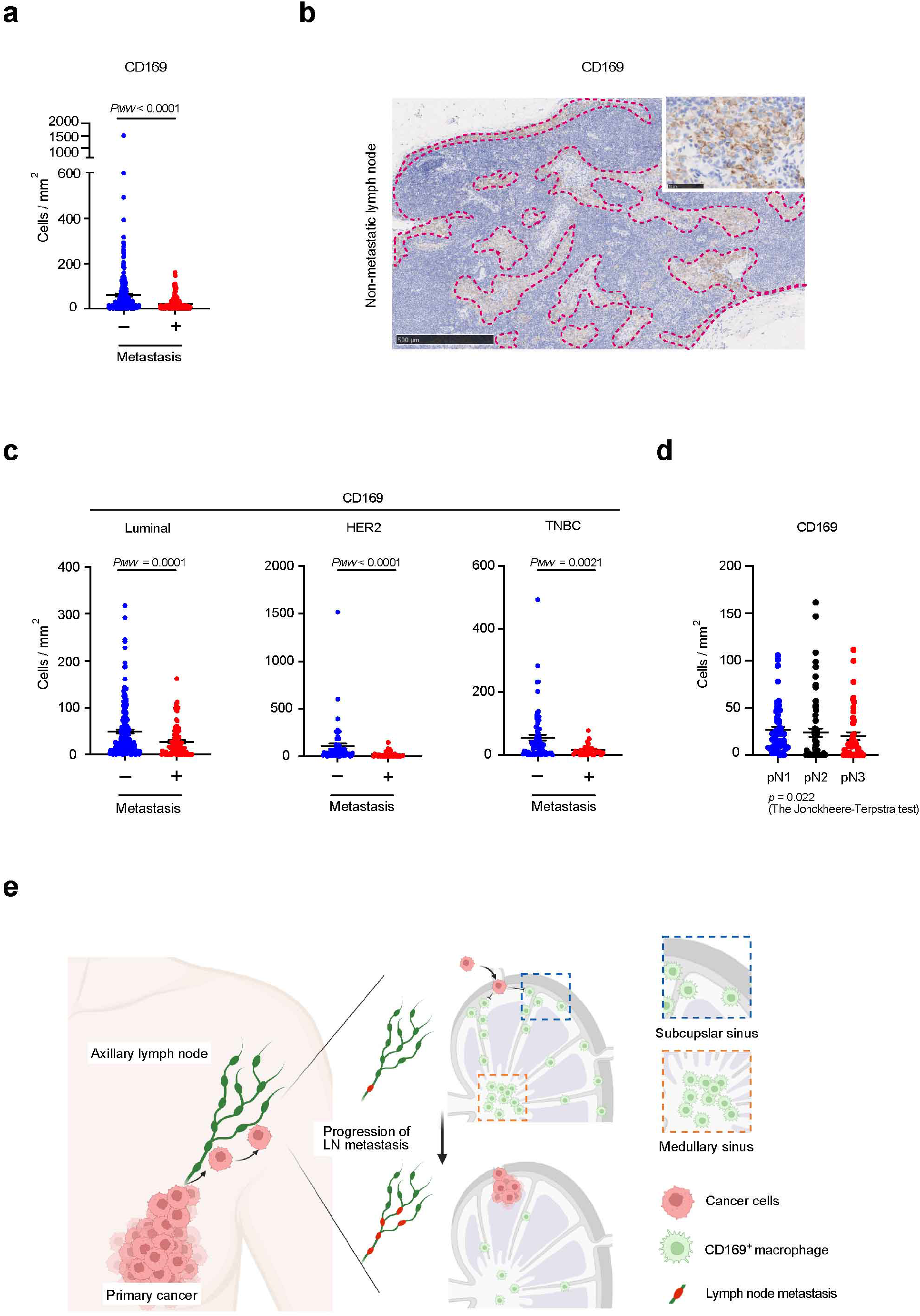
The elimination of CD169^+^ macrophages is a generalizable event in breast cancer patients. **a.** Density of CD169^+^ macrophages in non-metastatic and metastatic lymph nodes. Data are presented as the mean ± SEM. The *p* values were calculated using the two-tailed Mann-Whitney *U* test. A total of 315 non-metastatic and 159 metastatic lymph nodes were obtained from 58 patients. **b**. Representative immunohistochemical images of CD169 staining in non-metastatic lymph nodes. Dotted lines indicate the lymph node sinus. **c**. The density of CD169^+^ macrophages in non-metastatic and metastatic lymph nodes from luminal, HER2-positive, and triple-negative breast cancers. Data are presented as the mean ± SEM. The *p* values were calculated with the two-tailed Mann-Whitney *U* test. A total of 180 non-metastatic and 100 metastatic lymph nodes were obtained from 36 patients with luminal breast cancer; 59 non-metastatic and 32 metastatic lymph nodes were obtained from 10 patients with HER2-positive breast cancer; and 76 non-metastatic and 27 metastatic lymph nodes were obtained from 12 patients with triple-negative breast cancer (TNBC). See also Extended Data Table 5 for patient characteristics. **d.** The density of CD169^+^ macrophages in metastatic lymph nodes and pathological N classification. pN1 refers to 1–3 metastatic lymph nodes, pN2 refers to 4–9 metastatic lymph nodes, and pN3 refers to ≥ 10 metastatic lymph nodes. Data are presented as the mean ± SEM. The *p* values were calculated with the Jonckheere–Terpstra test. A total of 51, 62, and 46 metastatic lymph nodes were identified from 29, 18, and 11 patients with pN1, pN2, and pN3 disease, respectively. **e.** Graphical summary of the proposed mechanism. LN, lymph node.

We next investigated the correlation between the number of CD169^+^ macrophages and pN classification for breast cancer staging (pN1 = 1–3 metastatic lymph nodes; pN2 = 4– 9 metastatic lymph nodes; and pN3 ≥ 10 metastatic lymph nodes)^41^. Our data demonstrated a gradual decrease in CD169^+^ macrophages correlating with the pN classification (Fig. 5d). These results suggested that when a patient has more metastatic lymph nodes, more CD169^+^ macrophages are eliminated from each metastatic lymph node. However, pathological tumor size classification (pT) and volume of the metastasized cancer did not correlate with the number of CD169^+^ macrophages in the lymph nodes (Extended Data Fig. 5a,b). Furthermore, our slides included metastatic lymph nodes in which CD169^+^ macrophages accumulated in metastasized cancer tissues, despite being rare (15 of 159 investigated lymph nodes and 7.67 ± 14.1 cells/tumor in the 15 lymph nodes; Extended Data Fig. 5c). This was in contrast to our spatial transcriptome analyses, in which CD169^+^ macrophages were observed to be completely depleted around cancer tissues (Fig. 3e). These differences may reflect which stage of cancer-dependent suppression of CD169^+^ macrophages was captured in each lymph node section. In summary, our data strongly indicated that CD169^+^ macrophages were consistently eliminated from the metastatic lymph nodes of breast cancer patients, and that our experiments detected various different phases of CD169^+^ macrophage elimination (Fig. 5e).

## Discussion

The effective suppression of host anti-cancer immunity is vital for cancer cells to survive in the host environment. Cancer suppresses various immune cell types in the lymph node via remote, proximal, and direct mechanisms^17, 30, 33, 34^. The list of immune cell types suppressed by cancers in the lymph nodes remains incomplete. Moreover, in many cases, the precise mechanisms through which cancer suppresses immune cells and the invasion stage in which cancer cells achieve this suppression are unclear.

The current study establishes CD169^+^ macrophages as a target for proximal and/or direct suppression by cancers in the lymph nodes. Other lymph node cell types suppressed in cancer patients and mouse cancer models include CD4^+^ T cells, CD8^+^ T cells, and innate immune cells^3, 17–19, 30, 33, 34, 42–48^. T_reg_ cells are an important cell type involved in the suppression of CD4^+^ and CD8^+^ T cells in the lymph nodes^18, 19, 33, 34^. The elimination of CD169^+^ macrophages we identified in this study can be an additional mechanism leading to inability of T cells to combat cancer cells. In addition, cancer cells can remotely condition lymphatic vessels (i.e., non-immune cell type) to modulate the immune system via e.g., C-C motif chemokine 5 (CCL5)^30–32, 49^. Such conditioning may contribute to the elimination of CD169^+^ macrophages from the lymph nodes. Testing this possibility requires additional studies that compare lymph nodes from e.g., non-invasive breast cancer patients and non-metastatic lymph nodes from patients having lymph node metastasis.

We identified CD169^+^ macrophages as “early” targets of cancer cells in two meanings. First, CD169^+^ macrophages phagocytose cancer-derived antigens and present them to CD8^+^ T cells^20, 23^. This process represents an early step in anti-cancer immunity, the failure of which causes the corruption of the entire anti-cancer immunity^20–22^. Metastasized breast cancers inhibit this early process of anti-cancer immunity. Second, our data indicate that the elimination of CD169^+^ macrophages precedes other reported immune cell abnormalities, such as abnormalities in the altered number of immune cells^18, 19, 30, 42^ and systemic inflammation defined by CRP (Extended Data Table 4)^39, 40^. Thus, there is a possibility that the elimination of CD169^+^ macrophages might be an early job of metastasized cancer cells to achieve sufficient immunosuppression. Given the critical role of CD169^+^ macrophages in anti-cancer immunity^20–23^, their prioritized elimination appears to be reasonable for cancer cells, leading to the next important question of the underlying mechanism exploited by cancer cells to achieve this immune-evading effect.

The mechanism underlying the elimination of CD169^+^ macrophages from the lymph nodes is currently unclear, which is a major limitation of this study. However, our datasets successfully captured different stages of CD169^+^ macrophage suppression in patients (Fig. 5 and Extended Data Fig. 5c). According to these results, we propose a phased mechanism (i.e., sequence) of CD169^+^ macrophage suppression in the lymph nodes of breast cancer patients (Fig. 5e). Antigens from primary cancer tissues transported via the lymph may activate CD169^+^ macrophages, triggering anti-cancer immunity^20^. Cancer cells somehow break this defense, invading lymph nodes. Upon cancer cell arrival, CD169^+^ macrophages show some anti-cancer activity against metastasized cancer cells (Extended Data Fig. 5c). However, metastasized cancer cells can directly and/or indirectly expel CD169^+^ macrophages from the lymph nodes, achieving down-regulation of anti-cancer immunity. Such immunosuppression may allow expansion of metastasized breast cancer cells in the lymph nodes (Fig. 5d) and may cause other adverse effects including lowered efficiency of anti-cancer immunotherapy. Addressing the exact mechanisms behind CD169^+^ macrophage suppression in the lymph nodes is our next challenge.

We also suggest that the conflict between breast cancer cells and CD169^+^ macrophages in the lymph nodes is of significant pathophysiological importance. This is strongly supported by the correlation between breast cancer staging and the number of CD169^+^ macrophages in the metastatic lymph nodes (Fig. 5d). To the best of our knowledge, the number of CD169^+^ macrophages in metastatic lymph nodes is one of the strongest cellular indicators of breast cancer progression in patients, further highlighting the clinical implications of our discovery. Given these implications, we envision that retaining CD169^+^ macrophages in the lymph nodes in patients may be a way to increase the efficiency of anti-cancer immunotherapy in the future.

In summary, the current study identified suppression of CD169^+^ macrophages as the most prominent pathological phenotype in lymph nodes with breast cancer metastasis, establishing this suppression as a critical therapeutic target in the future.

## Methods

### Clinical samples and tissue processing

Lymph nodes were collected from six patients with breast cancer who underwent axillary dissection at Kyoto University Hospital (Kyoto, Japan) under the institutional ethical guidelines and regulations/ethical principles of the Declaration of Helsinki. The study protocol was approved by the Institutional Ethical Committee (G0424-18) (Kyoto University Graduate School and Faculty of Medicine) and informed consent was obtained. The clinical and pathological characteristics of the patients are summarized in Table 1. Lymph nodes were collected in the surgical room and cut in half. Enlarged lymph nodes were cut into one-quarter or one-eighth sections. The numbers of lymph nodes collected from each patient are summarized in Table 1. The collected tissues were embedded in OCT compound (Sakura Finetek Japan), frozen in liquid nitrogen, and immediately stored at −80°C. Tissue blocks were sectioned at 15 μm thickness using a Leica CM1950 cryostat microtome. Each section was mounted on polyethylene naphthalate membrane 4.0 μm slides (Leica). The sections were stained with hematoxylin and eosin (HE). Among the metastatic lymph nodes, only those with isolated metastatic masses were selected to prevent cancer cell contamination. Patients with lymph nodes exhibiting diffuse cancerous infiltration were excluded from the study. Lymphocyte regions were isolated using an LMD7000 laser micro-dissection system (Leica Microsystems) following the manufacturer’s protocol.

### Bulk RNA-sequencing

RNA was extracted using the RNeasy Micro Kit (QIAGEN) following the manufacturer’s protocol. Library preparation and sequencing were performed by Macrogen (Japan). Libraries were prepared using the SMART-Seq v4 Ultra Low Input RNA Kit and TrueSeq RNA Sample Prep Kit v2 following the manufacturer protocols (Library protocol SMART-Seq v4 Ultra Low Input RNA). Paired-end sequencing was performed using the Illumina NovaSeq 6000 platform with a 100-bp read length.

### Transcriptome analysis

RNA-sequencing reads were mapped to the human reference genome h38/GRCh38 using Hisat2^50, 51^ and counted using Feature Count^52^ in Galaxy (https://usegalaxy.org/). Read counts were normalized using the transcripts per kilobase million (TPM) method, and the low-expression genes (TPM < 1) and non-protein-coding genes were removed. Gene expression matrices generated using the TPM scores are listed in Extended Data Table 1.

### Spatial transcriptomics

Formalin-fixed, paraffin-embedded (FFPE) samples were used for spatial transcriptomic analysis. Tissue slides were prepared using Visium CysAssist Spatial Gene Expression for FFPE, following the manufacturer’s protocol (CG000518, 10x Genomics). Deparaffinization, HE staining, and de-crosslinking were performed following the manufacturer’s protocol (CG000520, 10x Genomics). Libraries for Visium were prepared following the Visium Spatial Gene Expression User Guide (CG000495, 10x Genomics). Paired-end sequencing was performed using a NovaSeq 6000 system (Illumina), following the manufacturer’s protocol.

### Spatial transcriptomics data analysis

Data processing: Spatial transcriptomics data, including UMI counts and spot coordinates, were analyzed using the R Seurat package (version 4.3.0)^53^. The four images and their data were processed into Seurat objects using the Load10X_Spatial function and normalized using the NormalizeData function. Visual inspection revealed that each sample contained spots with low numbers of reads and detected genes or spots detached from the main part of the tissue slice. Spots were manually filtered from each slice. Ultimately, 12,625 spots remained. The data of the four samples were merged, and batch effects among the four samples were removed using Harmony (version 0.1.1, function RunHarmony)^54^. The first 10 Harmony dimensions were used to conduct further dimensionality reduction (with the RunUMAP function) and spot clustering (with the FindNeighbors and FindClusters functions; resolution = 0.5) into 11 clusters. To interpret the cell-type composition of each spot, we inspected the expression patterns of known marker genes (Extended Data Table 2).

Analysis of biological pathways: We defined sets of genes associated with Gene Ontology (GO) biological process functional annotations using the R package msigdbr (version 7.5.1). We filtered out GO terms associated with <20 or >250 genes present in the data. For each of the remaining 3,152 GO terms and their associated genes (Extended Data Table 2), we calculated module scores using Seurat’s AddModuleScore function in the spots of the spatial transcriptomic data^53^. In brief, module scores reflected the average expression of the set of genes within each spot or cell subtracted from the average expression of several control genes with similar average expression levels as the input set. AddModuleScore was used with default parameters. In addition, within each of the 11 spot clusters, we predicted differentially expressed genes between lymph nodes with and without metastasis (Function FindMarkers). We performed this analysis for samples from each patient separately to avoid patient-specific biases.

### Antibody panel design

Information on the antibodies, metals, and their concentrations is provided in Extended Data Table 3. Unlabeled antibodies were conjugated with metals using the Maxpar X8 Multimetal Labeling Kit-40 Rxn (Standard BioTools) following the manufacturer’s protocol.

### Imaging mass cytometry

FFPE tissue samples were prepared at Kyoto University Hospital in Japan. FFPE section staining follows by modified manufacturer’s protocol (Fludigm, PN 400322 A3). The slides were baked for >2 h until all visible wax was removed. The slides were then deparaffinized in fresh xylene and ethanol. The slides were then inserted into preheated antigen retrieval ethylenediaminetetraacetic acid (EDTA) buffer (pH 9.0) and incubated for 30 min. Following incubation, the slides were cooled to 70L in the antigen retrieval EDTA buffer. The slides were washed twice with distilled water for 5 min each, followed by washing in phosphate-buffered saline (PBS) for 10 min each. The slides were blocked with 0.5% bovine serum albumin/PBS for 45 min at room temperature and incubated overnight with the antibody cocktail at 4℃ in a hydration chamber. The slides were washed twice with 0.2% Triton X-100 in PBS for 5 min and twice with PBS for 5 min each. The tissue was stained with the intercalator Ir in PBS (1:400) for 30 min at room temperature, followed by washing with distilled water for 5 min. The slides were air-dried for 20 min at room temperature. Imaging mass cytometry was performed using a Hyperion Imaging System (Standard BioTools). Regions of interest (ROIs) were selected from at least three points, including the nearest part of the invading tumor. The ROI was selected at 1000 μm width and 1000 μm height. Imaging was performed according to the manufacturer’s protocol at a laser frequency of 200 Hz and power of 3 dB.

Images were exported as TIFF files (OME-TIFF 32-bit) in an MCD viewer (Standard BioTools) and loaded into HALO version 3.5.3577 (Indica Labs). Imaging analysis was performed using the HighPlex FL module (v4.2.5) following the manufacturer’s protocol. Positive thresholds for individual markers were determined according to published nuclear or cytoplasmic staining patterns.

### Immunostaining and cell count

FFPE sections from the lymph nodes of patients with breast cancer were obtained from Kyoto University Hospital (Kyoto, Japan) according to the protocol approved by the Institutional Ethical Committee (R2889-1) (Kyoto University Graduate School and Faculty of Medicine). Immunohistochemistry for CD169 (sc-53442; clone HSn7D2; Santa Cruz Biotechnology, Santa Cruz, CA, diluted 1:25) was performed using the Ventana Discovery Ultra platform (Roche, Basel, Switzerland). Antigen retrieval was performed using the CC1 reagent (Ventana) for 60 min. The primary antibody was incubated for 32 min, and detection was performed using UltraView Universal DAB Detection Kit (Ventana). CD169-positive staining was counted in whole sliced lymph nodes on the slide, excluding lymph nodes completely occupied by cancer, by an experienced pathologist blinded to patient information.

### Statistical analysis and data visualization

The sample size was determined based on feasibility. Statistical significance was determined using a paired t-test for paired samples and the Mann–Whitney *U* test to compare cell count data between groups. Statistical analyses were performed using GraphPad Prism Software (Prism9). The Jonckheere–Terpstra test was used to evaluate the CD169 expression trend in cells according to pN and pT (SAS for Windows release 9.4; SAS Institute Inc., Cary, NC, USA). Statistical significance was determined at α = 0.05.

## Data availability

The bulk RNA transcriptome datasets used in this study are available in DNA Databank of Japan (DDBJ) under the accession numbers of DRR483580-DRR483599. The Visium datasets used in this study are available under accession number in the JGAS000616.

## Acknowledgments

This work was supported by JSPS KAKENHI (16H06279, 20H03451, 20H04842, and 22H04925; S.K.: 19K16770 and 21K15530; K.K.), Grant-in-Aid for JSPS Fellows (JP22KJ1822; Y.M.), AMED (JP21ck0106698; K.K.), JST FOREST (JPMJFR2062; S.K.), Caravel, Co., Ltd (S.K.), and Japan Foundation for Applied Enzymology (S.K.). This work was also partially supported by Sumitomo Pharma Co., Ltd. under SKIPS (Sumitomo-Kyoto University Innovation Promotion System). We thank the Center for Anatomical, Pathological, and Forensic Medical Research, Kyoto University Graduate School of Medicine, for preparing the microscope slides and performing immunohistochemistry. We also thank Sunao Tanaka, Keiko Muta, Pu Fengling, and Kayoko Koishihara for helping collect patient samples.

## Author contributions

K.K. conceived and supervised the study, provided patient samples and clinical information, designed experiments, analyzed data, constructed figures, and revised and edited the manuscript. S.K. supervised the study, designed experiments, analyzed data, constructed figures, and wrote the manuscript. Y.M. collected patients’ samples, performed experiments, analyzed data, constructed figures, and wrote the manuscript. T.R.K. provided patient samples, contributed to the experimental design, and performed the immunohistochemistry evaluation. A.V. performed spatial transcriptome analyses and constructed figures. M.H. prepared tissue samples for spatial transcriptome experiments. Y.S. performed spatial transcriptome experiments. Y.T. and H.H. provided patient samples and contributed to the experimental design. Y.F. collected patients’ samples and performed experiments. Y.I. and S.M. conducted the statistical analysis. M.T. conceived and supervised the study. All authors provided intellectual and reviewed the paper.

---

**Extended Data Fig. 1.**
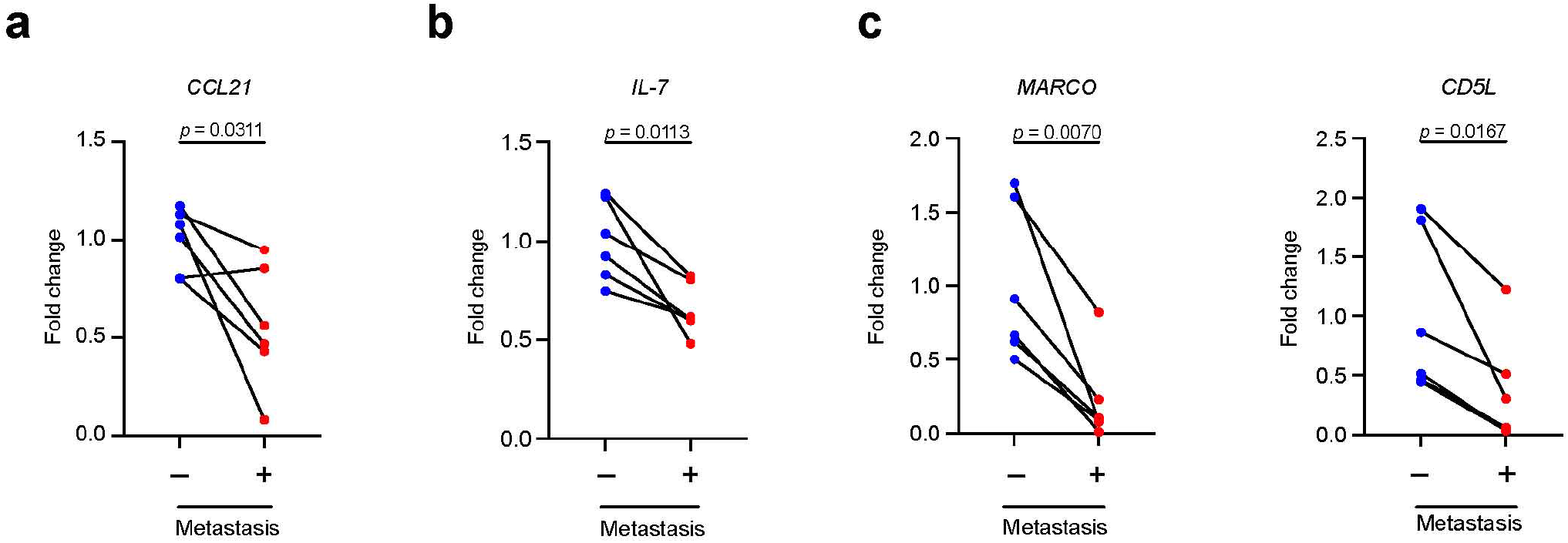
Representative genes down-regulated in metastatic lymph nodes. **a.** Expression of *CCL21*. **b**. Expression of *IL-7.* **c.** Expression of *MARCO* and *CD5L.* Averaged fold-change data (RNA-seq) normalized to non-metastatic lymph nodes are presented. *p* values were calculated with the paired two-tailed Student *t*-test (*n* = 6).

**Extended Data Fig. 2.**
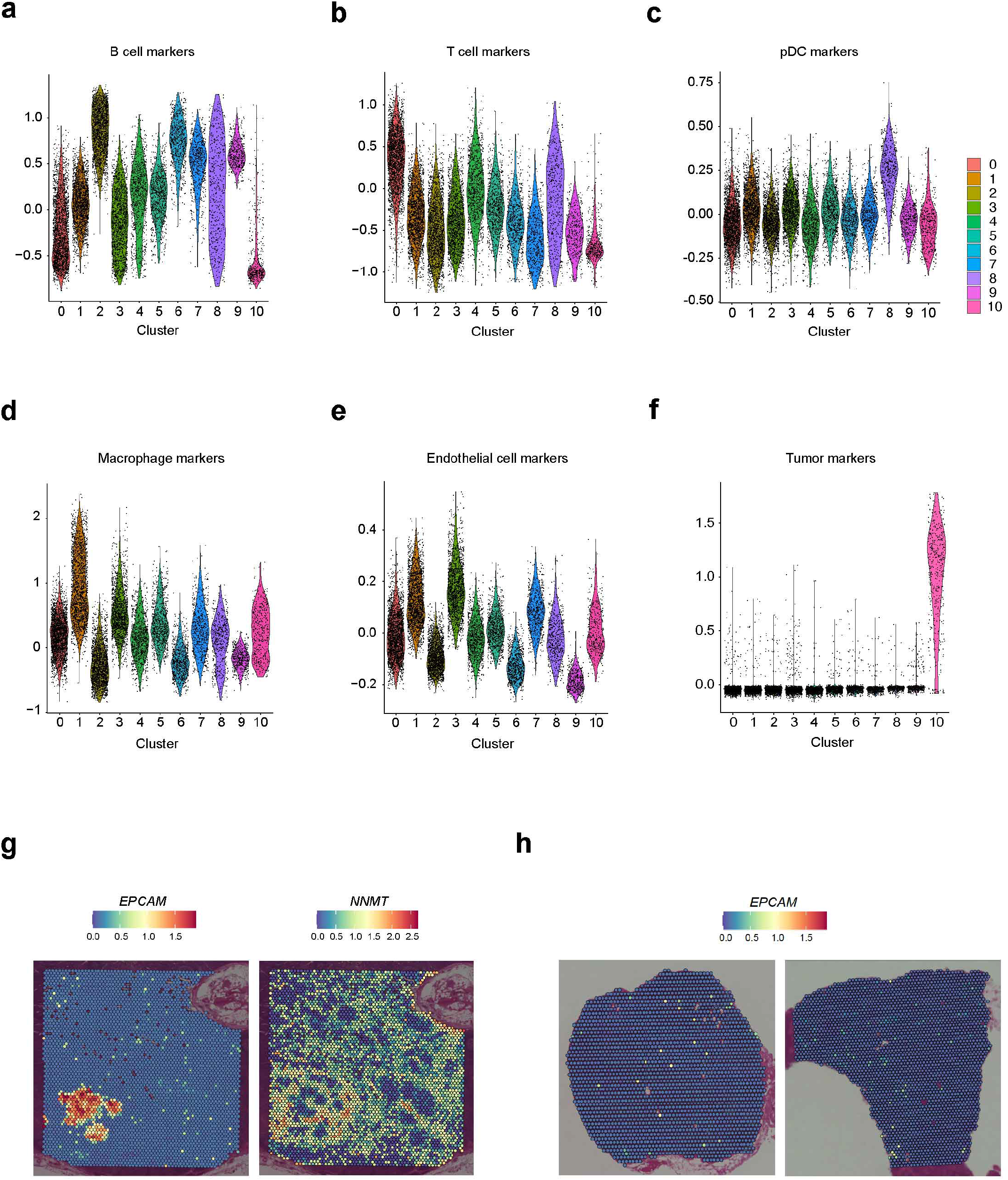
Characterization of VISIUM datasets. **a–f.** Violin plots for (a) B cell, (b) T cell, (c) plasmacytoid dendric cell (pDC), (d) macrophage, (e) endothelial cell, and (f) tumor markers in UMAP clusters. See Extended Data Table 2 for the complete list of marker genes used in this study. **g**. Expression of *EPCAM* and *NNMT* in metastatic lymph nodes. The *EPCAM* data are identical to those in Fig. 2e and are displayed again for direct comparison with *NNMT* expression. **h**. Expression of *EPCAM* in non-metastatic lymph nodes. No histologically visible metastases were observed in these two sections.

**Extended Data Fig. 3.**
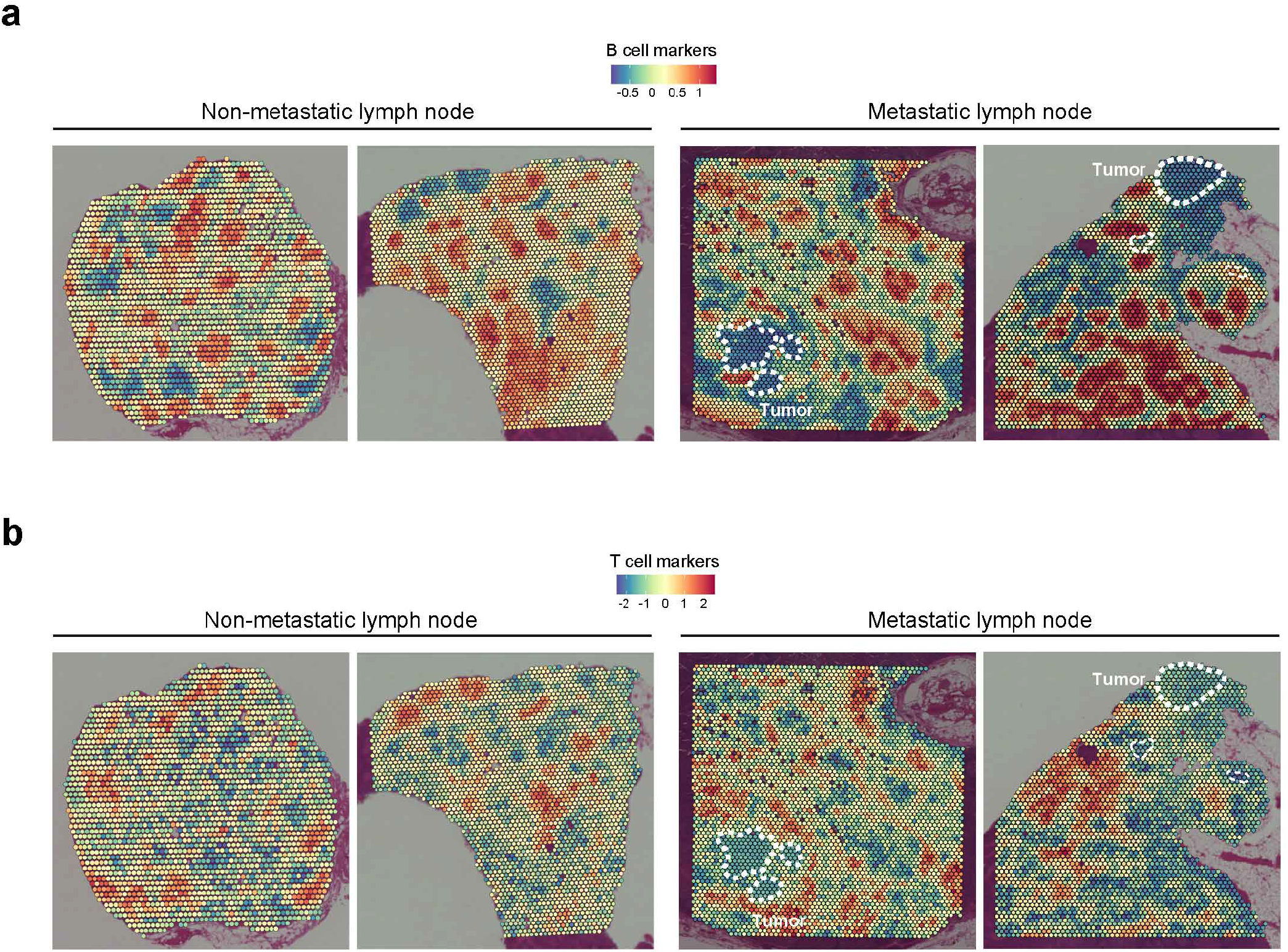
B- and T cell-enriched spots in VISIUM slices. **a.** B cell-enriched spots in non-metastatic and metastatic lymph nodes. Metastatic breast cancer cells are outlined. **b.** T cell-enriched spots in non-metastatic and metastatic lymph nodes. Metastatic breast cancer cells are outlined.

**Extended Data Fig. 4.**
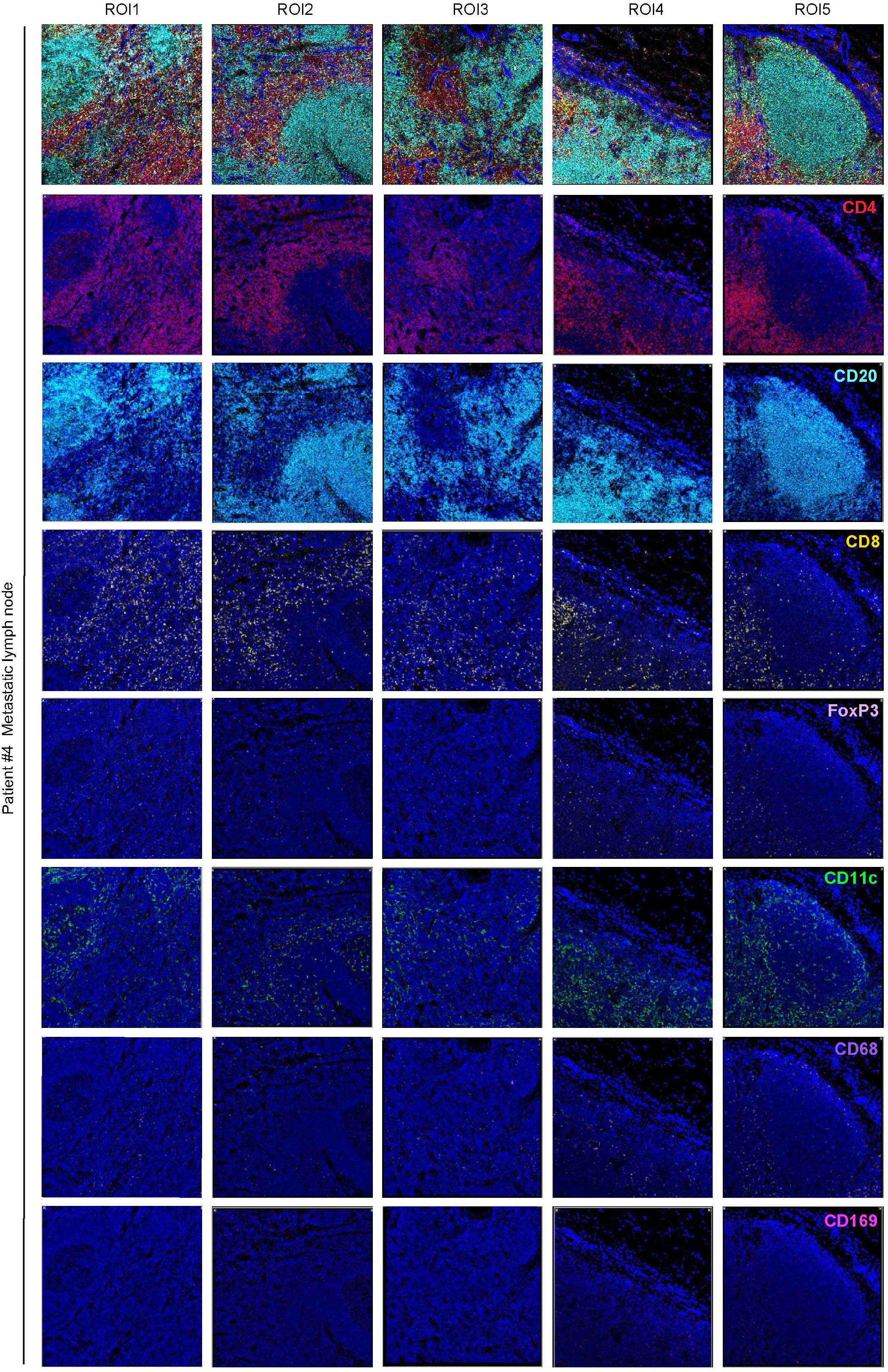
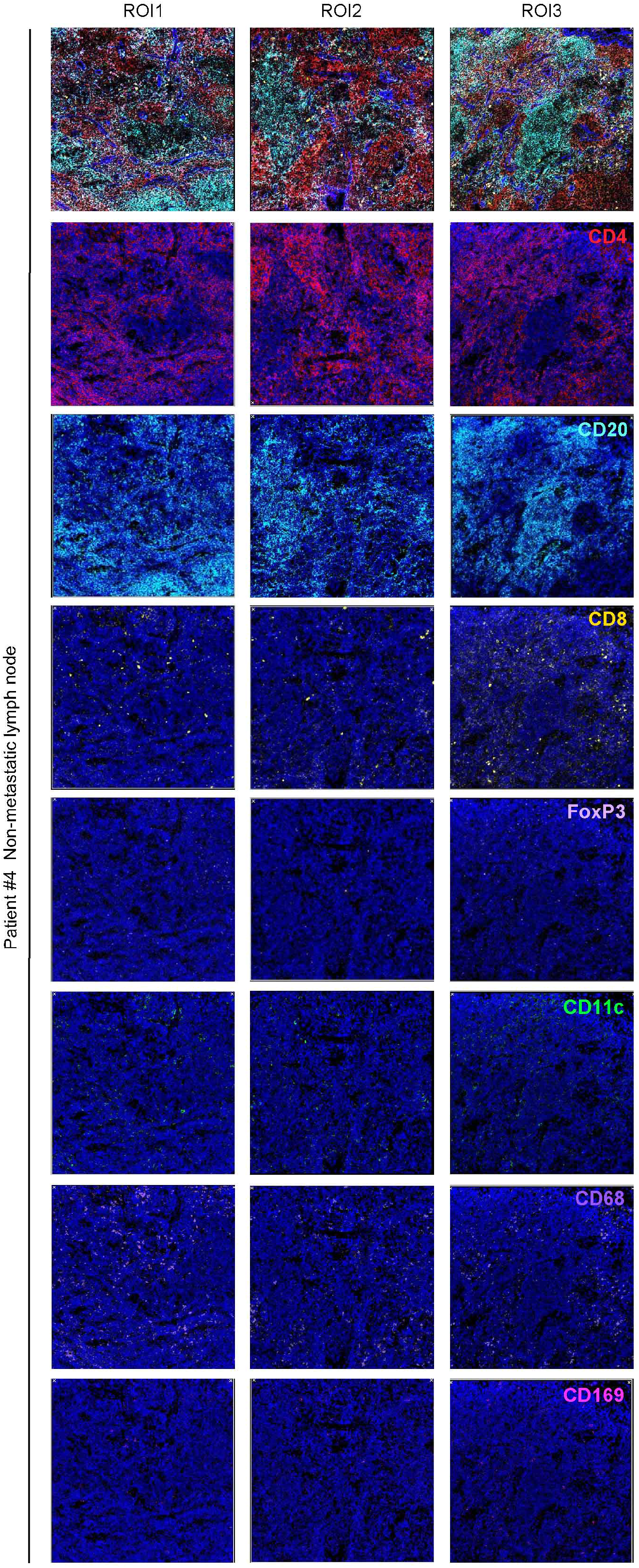
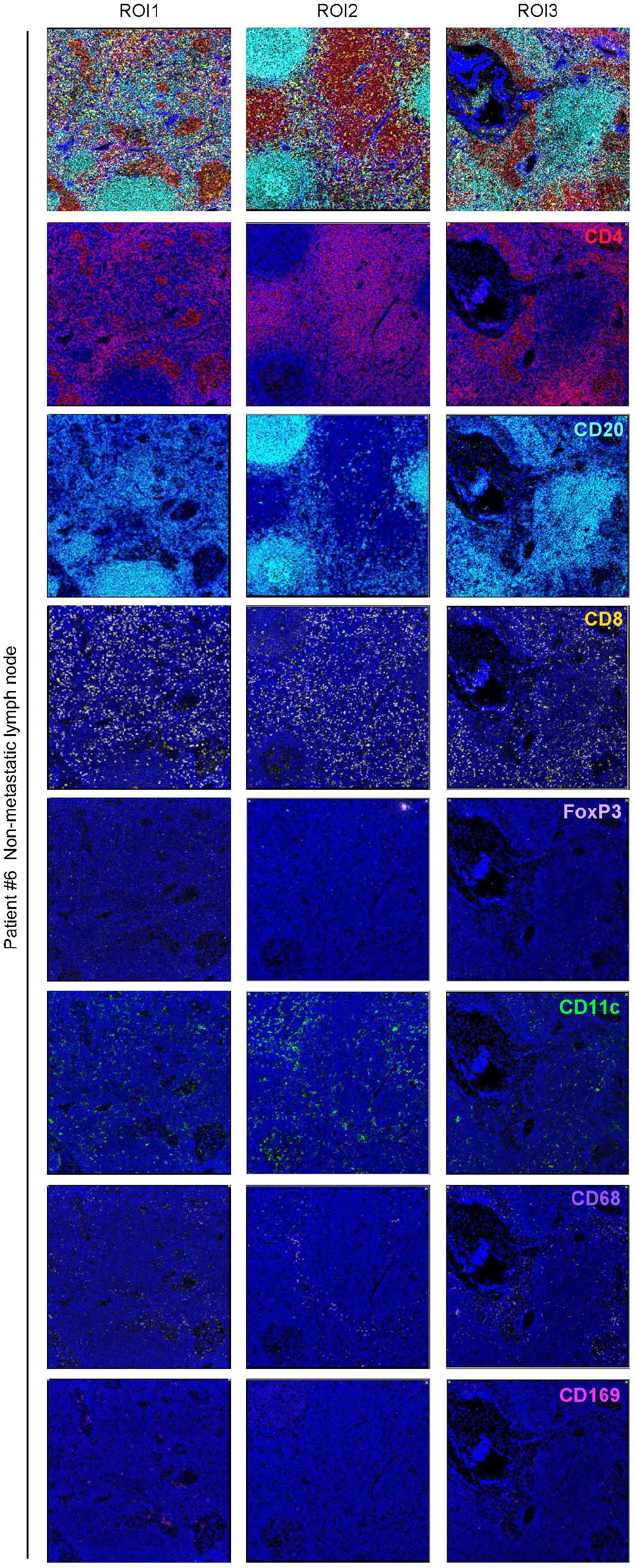
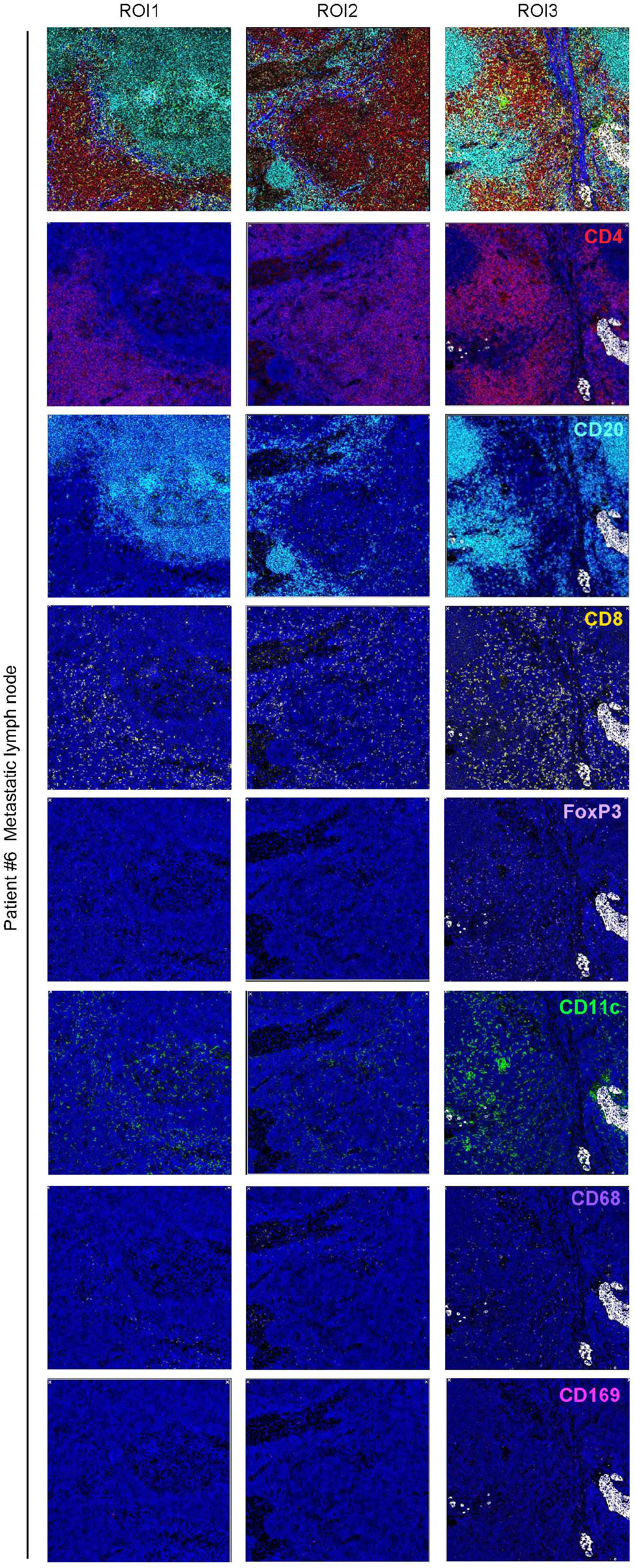
Raw pictures used for imaging mass cytometry analyses. Mass cytometry images of non-metastatic and metastatic lymph nodes from patients #4 and #6. At least three regions of interest (ROIs) are selected for each slice. Displayed channels are as follows: CD20 (B cells; cyan), CD4 (CD4^+^ T cells; red), CD8 (CD8^+^ T cells; yellow), FOXP3 (T_reg_ cells; pale purple), CD68 (macrophages; purple), CD169 (CD169^+^ macrophages; pink), CD11c (CD11c^+^ cells; green), pan-cytokeratin (cancer cells; white), DNA intercalator (nucleus; blue), and anti-α-smooth muscle actin (SMA) (myoepithelial cells; blue). DNA intercalators are displayed in each panel. The SMA staining is presented in the merged images. Cell phenotypes were determined using HALO software.

**Extended Data Fig. 5.**
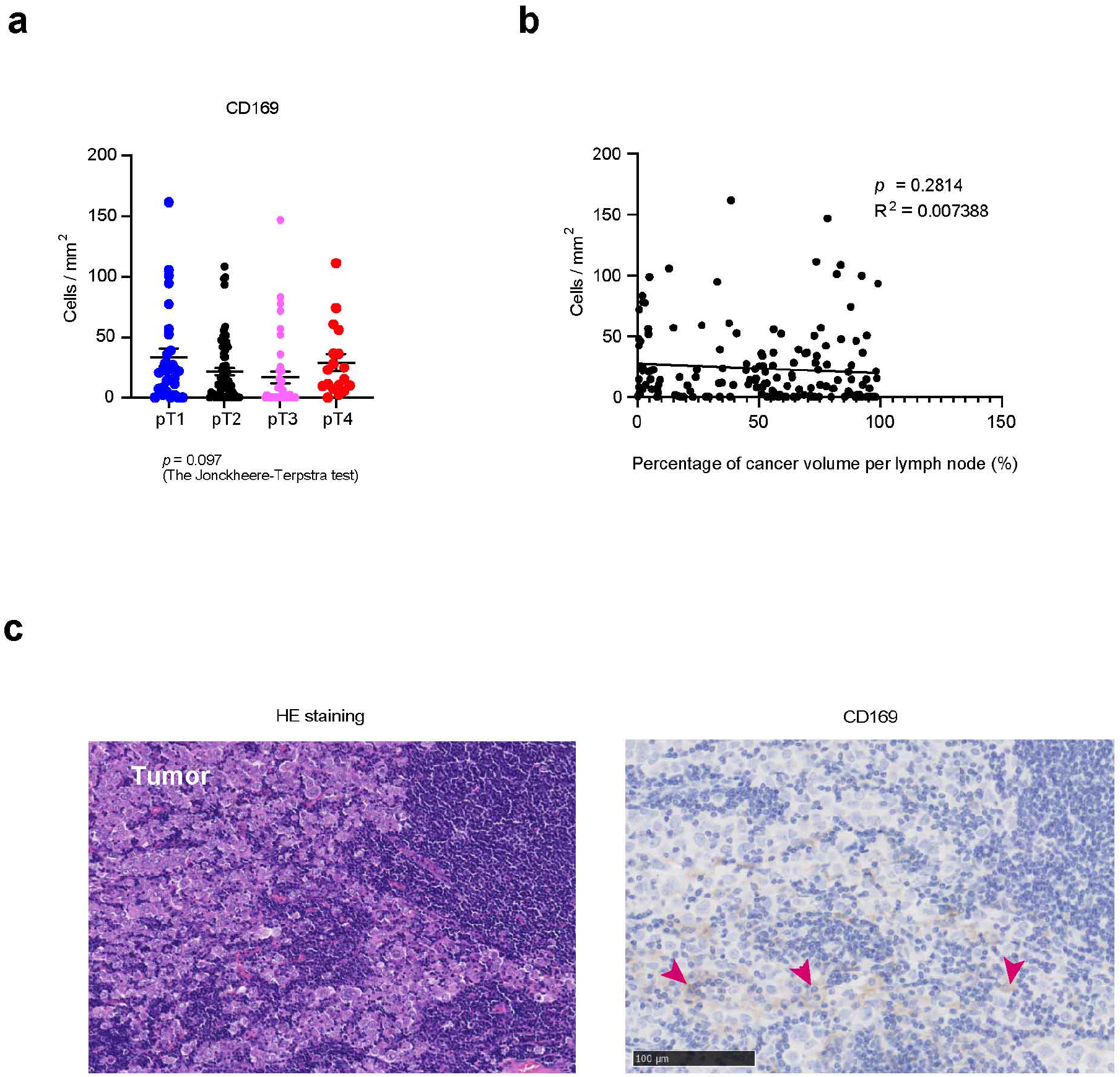
Relationship between clinical phenotypes and CD169^+^ macrophages. **a.** Associations of the density of CD169^+^ macrophages in the metastatic lymph nodes and pathological T classification. Data are presented as mean ± SEM. The *p*-value was calculated with the Jonckheere–Terpstra test. There were 30 metastatic lymph nodes in 14 patients with pT1 disease, 69 metastatic lymph nodes in 27 patients with pT2 disease, 42 metastatic lymph nodes in 11 patients with pT3 disease, and 18 metastatic lymph nodes in six patients with pT4 disease. **b**. Correlation between CD169^+^ macrophage density and metastatic cancer volume. The cancer volume was calculated as the percentage of the area in each lymph node section (*n* = 159). The *p*-value was calculated based on simple linear regression. **c.** Representative immunohistochemical images of CD169^+^ macrophages accumulating in metastatic cancer tissues. Left: hematoxylin-eosin (HE) staining; right: the immunohistochemical image of CD169 staining. Arrowheads indicate representative CD169^+^ macrophages within the cancer tissues.

